# Assembly and reasoning over semantic mappings at scale for biomedical data integration

**DOI:** 10.1101/2025.04.16.649126

**Authors:** Charles Tapley Hoyt, Klas Karis, Benjamin M. Gyori

## Abstract

**Motivation:** Hundreds of resources assign identifiers to biomedical concepts including genes, small molecules, biological processes, diseases, and cell types. Often, these resources overlap by assigning identifiers to the same or related concepts. This creates a data interoperability bottleneck, as integrating data sets and knowledge bases that use identifiers for the same concepts from different resources require such identifiers to be mapped to each other. However, available mappings are incomplete and fragmented across individual resources, motivating their large-scale integration.

**Results:** We developed SeMRA, a software tool that integrates mappings from multiple sources into a graph data structure. Using graph algorithms, it infers missing mappings implied by available ones while keeping track of provenance and confidence. This allows connecting identifier spaces between which direct mapping was previously not possible. SeMRA is customizable and takes a declarative specification as input describing sources to integrate with additional configuration parameters. We make available an aggregated mappings resource produced by SeMRA consisting of 43.4 million mappings from 127 sources that jointly cover identifiers from 445 ontologies and databases. We also describe benchmarks on specific use cases such as integrating mappings between resources cataloging diseases or cell types.

**Availability:** The code is available under the MIT license at https://github.com/biopragmatics/semra. The mappings database assembled by SeMRA is available at https://zenodo.org/records/15208251.

## 1 Introduction

Consistent and standardized identification of biomedical entities is an important step in creating and maintaining FAIR (Findable, Accessible, Interoperable, and Reusable) [56] data. However, many ontologies and databases that catalog biomedical entities are partially overlapping. For instance, cancer cell lines appear in several ontologies [4, 17, 47, 37], databases [5, 52, 14, 46], and generic nomenclature resources [45, 42, 7]. As a specific example, the human astrocytoma cancer cell line 13210N1 appears in Cellosaurus [4] under the identifier `cellosaurus:0110` and in the Cell Line Ontology (CLO) [47] as `CLO:0001072`.

Recognizing equivalent entries in different databases and mapping or merging across these is crucial for data interoperability and data analysis spanning multiple sources. It is also critical for computational tasks such as the construction of knowledge graphs [23, 11, 41], automated systems biology model assembly [19, 3], finding identifiers for bioentities mentioned in text [20], and integrating ontologies [13, 18, 35]. Each of these tasks requires mappings (also called *ontology mappings* or *semantic mappings*) across two or more resources that specify *equivalence* or other relations such as *broad, narrow*, or *related* between specific entries of the two resources. Mappings also ideally provide metadata on provenance and confidence to allow usage in a principled way. We refer to Figure 2 of [38] for a more detailed description of mappings. Overall, semantic mappings are crucial for consistent integration across resources.

### 1.1 Problem statement

Semantic mappings are often made available by individual (“primary”) resources, for example, most Open Biological and Biomedical Ontologies (OBO) [31] provide cross-references to equivalent or related entries in overlapping ontologies. In addition, “secondary” mappings resources including aggregators such as BridgeDB [50], TogoID [30], or independent mapping repositories like Biomappings [28] provide mappings that are collected from multiple sources or extend upon what is available from individual resources using additional curation. However, mappings remain difficult to assemble at scale because of the variety of *ad hoc* storage formats they are made available in, the ways they are produced (e.g., manual curation, rule-based inference, lexical matching), and the availability of metadata (e.g., precise mapping relations, curator confidence). A survey from [34] of mappings in life science ontologies highlighted several widespread issues such as the use of unspecific predicates (for example `oboInOwl:hasDbXRef` which is widely used to represent cross-reference mappings but is not specific enough to assert exact equivalence) and a lack of detailed provenance metadata. The Simple Standard for Sharing Ontological Mappings (SSSOM) [38] was developed to support curation of more specific predicates and detailed provenance metadata and is gaining adoption across many resources as a standard, however, it remains the case that most available mappings do not following a standard format or semantics.

Even if one were to assemble primary and secondary mappings at scale, there often exist gaps. **Figure 1** demonstrates that the landscape of mappings between cancer cell line resources is highly fragmented, and inference over multiple mappings is often necessary to identify key equivalences, e.g., between the BRENDA Tissue Ontology (BTO) [17] and CLO entities. The Ontology Mapping Service (OXO) [22] imports low-precision database cross-references from biomedical ontologies and implements inference across these, but is susceptible to data quality issues, explosions due to many-to-many mappings, and excludes valuable nomenclatures (e.g., the NCBI Entrez Gene Database [36]) that are not curated as ontologies. Notably, the landscape of mappings between resources is highly scattered, such that a combination of several mappings is required to identify equivalences, e.g., between the BTO entity and the CLO entity in Figure 1.

**Figure 1:**
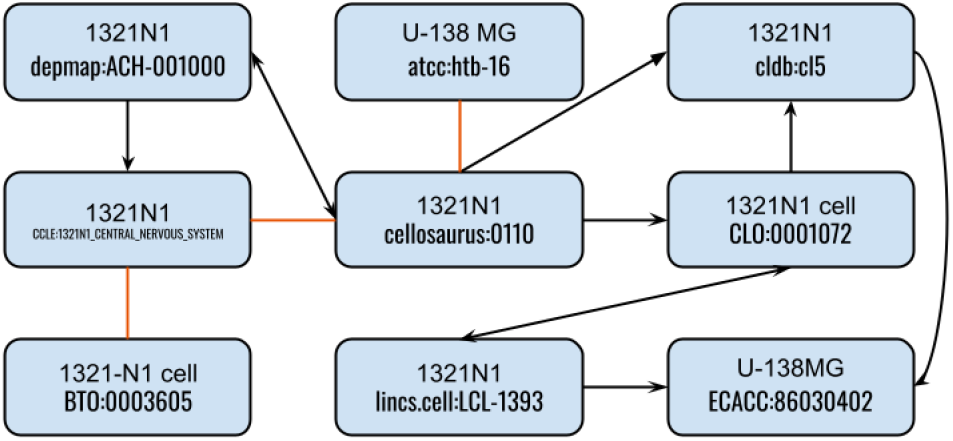
1321N1 is a cancer cell line derived from an astrocytoma of a 47 year old male caucasian patient. It is characterized by prominent pathogenic single nucleotide variants in the PTEN (`HGNC:9588`)) and TP53 (`HGNC:11998`)) genes. It appears in a variety of cell line resources (e.g., Cellosaurus), anatomy resources (e.g., BTO), clinical databases (e.g., DepMap, CCLE), and vendor catalogs (e.g., Sigma Aldrich). Solid black lines denote primary mappings pointing away from the resource from which the mapping was derived. Orange lines denote secondary mappings from Biomappings [28].

**Figure 2:**
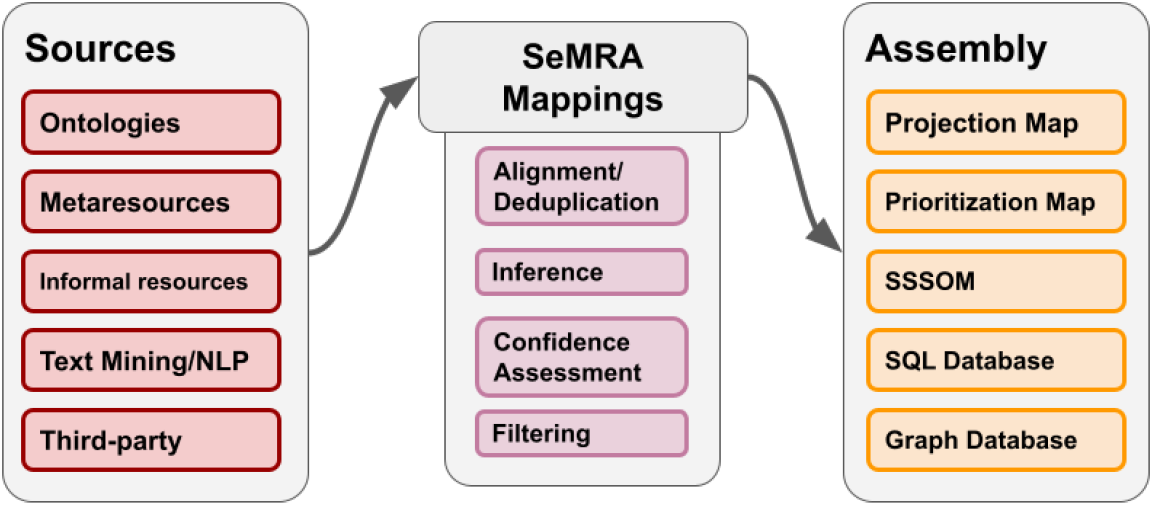
An overview of the software architecture of the SeMRA package. Source modules provide ingestion logic from various sources, feeding into a set of central SeMRA Mappings modules that compile mappings serving as input to Assembly modules that produce outputs in multiple export formats.

There are several other problems that currently limit the use of mappings for data integration. First, there does not yet exist a provenance model that captures the combination of mappings and algorithmic approaches used to derive an inferred mapping. Second, there do not yet exist methods that incorporate uncertainty associated with the provenance of a mapping into measures of confidence for inferred mappings. Uncertainty results from, for example, noisy mapping algorithms (e.g., lexical prediction), implicit one-to-many mappings due to broad or narrow matches, and curation error. An appropriate confidence model would incorporate both assumptions about how a mapping is produced and its intended interpretation for a specific data integration task. Third, there does not yet exist a data model nor workflow to declare implicit prior knowledge about the interpretation of cross-references from specific sources. For example, it’s known that the mappings distributed in an *ad hoc* TSV file from the Cancer Dependency Map Project (DepMap) [5] to Cancer Cell Line Encyclopedia (CCLE) [14] are *equivalences*, while only a subset of the low-precision mappings in Cellosaurus to CLO represent equivalences.

### 1.2 Contribution

To address these issues and open problems, we introduce the Semantic Mapping Reasoning Assembler (SeMRA), a novel method for automatically assembling mappings at scale, implemented as configurable open-source software. SeMRA represents mappings as a directed graph and provides functionality to infer indirect mappings based on graph traversal, then determine associated confidence. Importantly, it allows for customizing prioritization order to merge equivalent concepts during data integration in a consistent way. SeMRA further implements graph-based algorithms for flagging mappings that lead to inconsistent data integration as part of a semi-automated quality assurance workflow. It implements a web application for the interactive exploration of mappings, visualization of mapping graphs, and curation of flagged mappings.

We used SeMRA to assemble 43.4 million raw semantic mappings from 127 sources (including widely used biomedical ontologies and databases) to make available the broadest-coverage integrated mapping resource to date. In addition, we provide detailed analysis and metrics of using SeMRA on specific areas of biomedicine crucial for data integration. First, we demonstrate integrating cell and cell line resources using SeMRA, significantly expanding on results from [28]. Second, we integrate resources cataloging diseases and show substantial expansion in scope compared to results published by [21]. Finally, we provide metrics on the effect of semantic integration using SeMRA on overlaps between resources in five areas of biomedicine.

SeMRA is available as an open-source Python package (https://github.com/biopragmatics/semra) and provides a comprehensive mapping database in the SSSOM exchange format archived on Zenodo at https://zenodo.org/records/15208251.

## 2 Methods

SeMRA is built with a modular architecture with independent components for processing input sources, a semantic mapping data structure, inference and processing, assembly, export into multiple file formats, export to a graph database with a JSON API, and an interactive web application.

### 2.1 Sources

SeMRA implements a variety of import modules and processors for mappings. Where possible, SeMRA wraps preexisting parsers for standard representations. For instance, SeMRA reads mappings from ontologies in OBO format by wrapping the PyOBO Python package [29]. Similarly, SeMRA reads mappings from ontologies in the OWL and OBO Graph JSON formats using the Bioontologies Python package [25]. However, there are a number of ontologies that require custom processing due to nonstandard encoding of mappings, such as the Cell Line Ontology [47]. These are implemented in SeMRA with additional custom logic.

SeMRA also reads mappings in the SSSOM format via the `sssom-py` Python package [8]. This also gives access to several other formats, including the Expressive and Declarative Ontology Alignment Language (EDOAL; [9]) and the RDF Alignment Format [2], both used by the Ontology Alignment Initiative. Notably, support for SSSOM allows SeMRA to import mappings from Biomappings [28], a resource dedicated to predicting and curating mappings missing from primary sources.

SeMRA integrates with Wikidata [53] in order to automatically generate SPARQL queries for properties corresponding to various ontologies and databases (e.g., `P2158` for the Cell Line Ontology).

More generally, SeMRA implements a plug-and-play concept to implement custom parsers and importers for sources that are not in a standard format. Consequently, we implemented custom parsers to import mappings from OMIM, IntAct, PubChem, NCIT, ChEMBL, and several other resources. Similarly, SeMRA can be extended by users to process additional resources that are not available in a standard form.

### 2.2 Semantic mapping data structure

SeMRA implements a hierarchical data structure to represent mappings. The core data structure represents each unique mapping triple with a *subject, predicate*, and *object*. The *subject* and *object* represent entities in ontologies or databases using the compact universal resource identifier (CURIE) syntax [6]. CURIEs are automatically validated against the Bioregistry standard [27] to ensure consistent usage. The predicate comes from Simple Knowledge Organization System (SKOS) or related vocabularies (see https://mapping-commons.github.io/sssom/mapping-predicates/).

Each core triple has one or more evidence objects associated with it. There are two types of evidence objects. A *simple* evidence object contains a mapping justification based on the Semantic Mapping Vocabulary (SEMAPV) [39], an optional author annotation using an ORCID identifier, an optional confidence annotation from the producer, and a reference to the mapping set from which it came. Each mapping set has a name, version, license, and consumer confidence annotation. A *complex* evidence object represents the results of reasoning and contains pointers to all mappings used to construct the evidence, a mapping justification from SEMAPV, and an optional confidence factor reflecting the consumer’s confidence of the reasoning approach.

SeMRA’s mapping data structure is best suited for sequential operations. For other operations, including inference (see Section 2.3 below), a directed graph data structure is more appropriate. Each SeMRA edge can be compiled into a graph by making the subject of the mapping the source node, the object of the mapping the target node, and the predicate (and related evidences) properties on the edge. Mapping triples are inherently directional, despite many predicates being symmetric. Therefore, SeMRA explicitly models the directionality of each predicate while enabling inference of symmetric mappings in order to explicitly track provenance of operations.

#### Comparison to SSSOM

SeMRA’s data model captures all required and commonly used optional fields in SS-SOM. However, a conceptual distinction is that SSSOM represents individual mapping triples with a single supporting evidence as part of the mapping data structure, whereas SeMRA decomposes the representation of triples and the associated evidence. This allows for integration of evidence from multiple sources and a more sophisticated inference logic.

#### Confidence

Confidence associated with a mapping is calculated in a hierarchical manner based on the set of evidences supporting it, inspired by [3]. We interpret the confidence of a specific evidence for a mapping as a probability value between 0 and 1, and denote it *c*_*e*_ for a given evidence *e*.

To characterize *c*_*e*_, in case of a *simple* evidence, SeMRA takes into account both the producer’s confidence (if any provided) for its mappings *c*_Producer_ and the confidence of the consumer (in this case SeMRA or its user) in the mapping set the evidence came from: *c*_Mapping Set_. The evidence confidence is then expressed as *c*_*e*_ = 1 *−* (1 *− c*_Mapping Set_)(1 *− c*_Producer_). If no producer confidence is given, it is assumed to be 1.0, therefore simplifying the formula to directly using the consumer’s confidence in the mapping set. Assuming a mapping is supported by a set of evidences *E*, its overall confidence is expressed as the probability that at least one supporting evidence is correct: *c*_*m*_ = 1 *−*Π_*e ∈ E*_ 1 *− c*_*e*_ for mapping *m*.

Finally, the confidence for a *complex* evidence is calculated from the confidence of the mappings it is derived from: *c*_*e*_ = *γ*_*op*_(1 *−* Π_*m∈M*_ (1 *− c*_*m*_)) where *γ*_*op*_ is a scaling factor specific to reasoning operation *op* used.

#### Identifying mappings through hashing

Several applications require the unique identification of mappings and evidences associated with mappings. However, this is difficult because each data structure composes multiple entities. In SeMRA’s context a mapping’s identifier has to reflect the content of the mapping it represents to avoid inconsistencies in the highly dynamic and decentralized nature of the creation of mappings and evidences. SeMRA therefore implements a deterministic method for hashing mappings, evidences, and mapping sets independently that can be used for unique identification of each resource within the SeMRA ecosystem.

### 2.3 Inference and processing

SeMRA implements three inference methods: inversion, mutation, and transitivity. Using these inference methods, new mappings can be inferred from existing mappings or existing mappings can be made more specific, thereby broadening downstream data integration capabilities. These inference methods implemented in SeMRA are consistent with chaining rules described in the SSSOM documentation (**Figure 3**)^1^ to ensure consistency with community expectations of inferred mappings. These functionalities are exposed along with several filtering approaches through a high-level function called `semra.pipeline.process()`. We next describe each inference method in detail.

**Figure 3:**
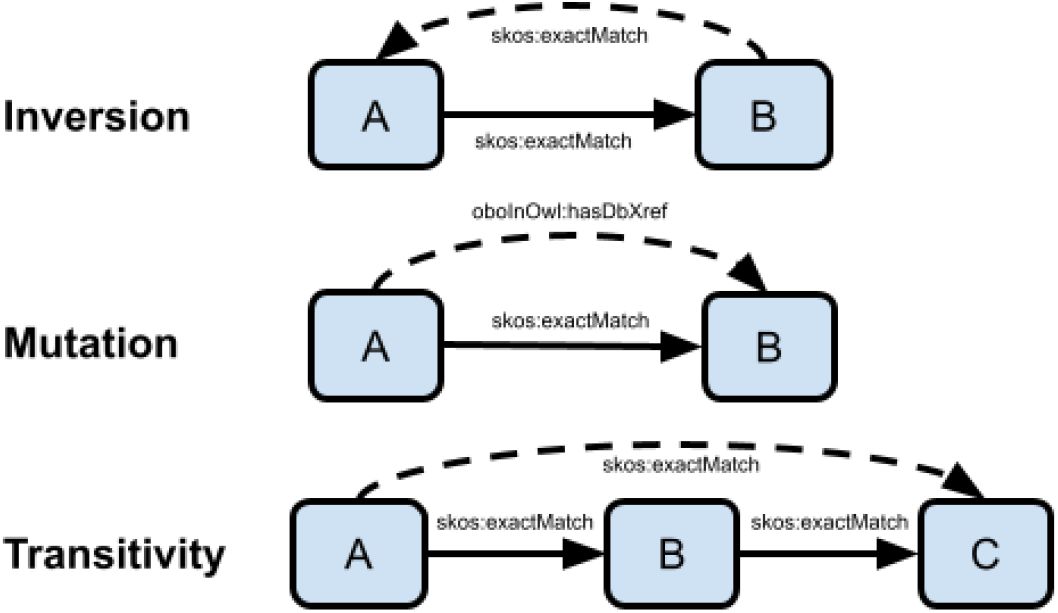
An overview of inference methods implemented in SeMRA, nodes representing entities between which edges represent mappings. The edge labels show examples of predicates involved in the inference steps with solid lines representing original edges and dashed lines representing inferred edges.

#### Inversion

Inversion creates a new mapping where the subject and object are swapped and the predicate is changed. For symmetric predicates, such as those representing equivalence, the same predicate is reused. For example, if A is an exact match of B, then B is also an exact match of A. For asymmetric predicates, a correspondence between inverse predicates is maintained to facilitate inversion. For example, if A is a narrow match of B, then B is a broader match of A. In a directed mapping graph, this is equivalent to adding a reverse edge.

#### Mutation

Mutation creates a new mapping in which the original predicate is replaced by a new one. Mutations that generalize a stronger predicate with a weaker one are pre-configured in SeMRA. This is useful when integrating sources using a combination of strong OWL-based semantics and weaker SKOS-based semantics. For example, if A is an OWL equivalent class to B, then A is also a (SKOS) exact match to B. In a directed mapping graph, this is equivalent to adding a parallel edge.

Mutations that ascribe stronger semantics to a weaker predicate are also useful, but are often context- or task-specific. For example, some modeling approaches do not distinguish between genes and proteins, and therefore use their identifiers interchangeably. In this scenario, the “has gene product” (RO:0002205) relation between genes and proteins can be safely mutated to an exact match predicate. Similarly, many OBO Foundry ontologies store implicitly exact matches in the database cross-reference field, and therefore lose their precision. SeMRA allows the prior knowledge about each ontology’s curation practices to be explicitly configured during inference.

#### Transitivity

Transitivity rules allow new mappings to be inferred from chains of two or more mappings. For example, if A is an exact match to B and B is an exact match to C, then A is an exact match to C. Transitivity rules can also incorporate more complex chaining rules with different predicate types. Inference based on transitivity rules is implemented in SeMRA using the directed mapping graph data structure. With this, it implements a custom graph traversal related to breadth-first search that incorporates transitivity rules, resolution of conflicting mappings, and filtering of negative mappings. In a directed mapping graph, this is equivalent to adding an edge corresponding to a path through the graph.

#### Filtering

SeMRA implements several filtering operations, which are often useful to automatically identify and remove false positive mappings generated by inference. First, filtering mappings based on their source prefix, target prefix, or source-target prefix pair is often useful to remove irrelevant mappings for a given context. Similarly, some pairs of source-target prefixes can be excluded from prior knowledge. For example, terms in disease-related ontologies like MONDO [51] should never be mapped to terms in gene-related resources like the NCBI Entrez Gene Database [36]. Second, filtering by cardinality includes the identification of entities in one resource that have exact matches to more than one entities in another resource. These do not allow consistent data integration, and present an opportunity for additional curation. Third, filtering by confidence presents another automated method for removing potentially low confidence mappings produced by a combination of low confidence steps, such as mutation.

### 2.4 Assembly

SeMRA provides several high-level workflows for assembling mapping sets into useful artifacts, several input and output functions for reading and writing mappings, and several export modes for building mapping databases.

#### Projection mapping set

A projection mapping set contains mappings that support mapping identifiers from one ontology or database to another. In a projection mapping set, all subjects’ prefixes are the same, and all objects’ prefixes are the same, however, they may contain a combination of simple and complex evidences arising from primary sources and inference approaches. Projection mappings help data integration between two specific resources. SeMRA implements a workflow for creating a projection map that can be stored in either SSSOM or a highly compressed tabular format.

#### Prioritization mapping set

A prioritization mapping set defines for each entity what the standard representation of that entity should be. Therefore, each entity appears as the subject in exactly one mapping, and some entities may appear as the object of several mappings. Effectively, a prioritization mapping induces star graphs for each clique of mutually equivalent entities. In order to create a prioritization mapping set, a prioritization list of prefixes is required. There is also an assumption that only one entity from each prefix is in each clique.

A prioritization mapping set is useful in knowledge integration tasks where multiple sources with overlapping identifiers sources (such as some of the resources shown in Figure 1) are used, across which terms need to be standardized to a single prioritized identifier.

### 2.5 Export and exploration

SeMRA provides several export formats. First, it provides a low-level, direct export of its data using Python’s `pickle` module. Second, it provides an export to SSSOM, which enables conversion via `sssom-py` to several other formats (RDF, OWL), and provides an upload option to the NDEx platform [43].

#### Graph Database

SeMRA implements a workflow for serializing a mapping set into the Neo4j graph database (https://neo4j.com) which enables custom queries and can serve as a back-end for user applications. The graph represents concepts, evidences, mappings, and mappings sets as nodes. It also represents mappings as edges between two concepts (representing mappings both as nodes and edges provides query flexibility), and further represents as edges the interdependencies between mapping sets, evidence, mappings, and concepts. SeMRA exports files for building the database along with a Docker configuration and startup scripts to enable any mapping set to be turned into a database.

#### Web Application

SeMRA also implements a locally deployable web application on top of its graph database that exposes several high-level operations as a JSON API built with FastAPI. It implements a user interface containing a dashboard for summarizing the concepts, mappings, evidences, and mapping sets in the database as well as for exploring mappings, concepts, and cliques of equivalent entities (**Supplementary Figure S2**). The SeMRA web application has an optional integration with the Biomappings package [28] for the curation of negative mappings. It gives users direct insight into cliques that have problematic features such as containing multiple entities from the same resource that can be curated then incorporated into future inference.

## 3 Results

We first present an overall integrated database of mappings using SeMRA. Then, to show SeMRA’s configurability and use case-specific utility, we provide several processing pipelines with more specific scope, as follows. In Section 3.2, we integrate semantic mappings across ten cell type and cell line resources. In Section 3.3, we apply SeMRA to integrating resources that catalog human diseases to expand on the analysis in [21]. In Section 3.4, we quantify the effect of semantic integration with SeMRA on resolving redundancies in five areas of biomedicine (including cells and diseases from the previous two case studies).

### 3.1 Integrated mapping database

Using SeMRA, we generated a database of 43.4 million mappings from 127 sources, including ontologies, databases, and custom sources. Of these, 36.8 million were exact matches, 5.5 million were imprecise database cross-references, and around 1 million were other types of mappings. The mappings cover entities from 445 different ontologies and databases, due to the fact that a given source often maps to several external ontologies and databases. SeMRA imports life and natural science ontologies from the entirety of the OBO Foundry [31], a high-quality subset of BioPortal [55], and other ontologies that have been curated in the Bioregistry [27] based on community-driven use cases. SeMRA also imports databases that are generally of interest in biomedical data integration through ontology adapters in the PyOBO software package or through custom importers. For example, this includes widely used chemical databases like ChEMBL [12] and gene databases such as NCBI Entrez Gene [36].

The SeMRA Mapping Database is available from Zenodo at https://zenodo.org/records/15208251 as an SSSOM file and as a pre-built graph database deployable via Neo4j. The SSSOM format explicitly states the license for each mapping, ultimately reflecting the mapping source’s license.

This demonstrates the application of SeMRA at scale, to a general collection of widely used resources.

### 3.2 Landscape of cell and cell line resources

Resources cataloging cell types and specialized disease cell lines are a cornerstone of biology research and are crucial for the interpretation of single-cell data [33] and cancer gene dependencies [49]. Multiple resources exist providing entries for cell types and cell lines, including dedicated databases such as Cellosaurus and CCLE as well as resources with a broader scope such as the Experimental Factor Ontology [37], or UMLS [7]. The major challenge remains that these databases overlap, but mappings between them are limited.

We used SeMRA with a declarative configuration as input to run semantic integration on ten relevant resources (**Supplementary Table S1**). In addition to primary mappings sources from each resource, SeMRA made use of the semi-automatically curated mappings described by [28]. SeMRA inferred thousands of new mappings that would not have been available without such integration. This included the production of the first mappings between several resources, including EFO-UMLS, CCLE-CLO, MeSH-Cellosaurus, and BTO-NCIT which have thus far not been available from other sources.

Next, we used SeMRA to programmatically identify several systematic issues in cross-references from Cellosaurus, which often map to multiple terms in the Cell Line Ontology. We used the SeMRA web application to interactively review these issues, identify the primary mappings causing many-to-many references, and directly curate overrides in Biomappings. In several situations where database cross-references were to narrower or broader terms, we were able to curate negative mappings with the SeMRA web application’s Biomappings plugin. We found this was the case since there are often duplicate terms in CLO for the same cell line, imported from multiple external resources without normalization. This suggests that further work is necessary to address issues where a given resource has multiple terms for the same concept that should be mapped jointly. The SeMRA Cell and Cell Line Landscape is available as a set of data files and Dockerized graph database at https://zenodo.org/records/15164183.

### 3.3 Landscape of disease resources

The characterization and classification of diseases has a rich history. Computational resources cataloging diseases enable the standardized annotation of health data from clinical trials, drug indications, electronic health records, and other artifacts. They further support computational efforts in drug discovery, target prioritization, and modeling. However, the number of tasks they support has led to a high proliferation of disease resources with high overlap.

We used SeMRA with a declarative configuration as input to run semantic integration on 16 relevant resources (**Supplementary Table S2**) including ontologies, databases, billing code systems, and generic resources. Our analysis showed that there is high redundancy between these resources that requires a union and deduplication via a combination of mappings from primary resources, secondary resources, and inference. While MONDO and UMLS have each been previously positioned as disease mapping hubs, our analysis suggests that they are by themselves insufficient, and that an automated mapping assembly pipeline can serve as a generalization of the purpose of MONDO as a mapping hub suggested by [21].

Unlike the cell line scenario, where the described resources could be directly downloaded, the disease resource scenario presents the problem that several resources (ICD10-CM, IC9, ICD9-CM, and OMIM) can not be directly downloaded either due to lack of availability or usage restrictions (in the case of OMIM). Therefore, this analysis is only able to estimate a lower bound for the number of terms appearing in these resources based on the ones appearing in mappings. The SeMRA Disease Landscape is available as a set of data files and Dockerized graph database at https://zenodo.org/records/15164180.

### 3.4 Meta-landscape analysis

We performed landscape analyses over three additional domains to the first two case studies (anatomy, protein complexes, and genes) in order to estimate the total number of terms across common resources, as well as to use mappings to estimate the number of consolidated terms (**Table 1**). Configurations for each domain can be found at https://github.com/biopragmatics/semra/tree/main/src/semra/landscape.

**Table 1:** Meta-landscape analysis statistics. Each Zenodo record contains all raw mappings, processed mappings, and a Dockerfile for running the graph database, JSON API, and SeMRA web application for the domain.

| Domain | Raw Terms | Unique Terms | Reduction | Zenodo ID |
| --- | --- | --- | --- | --- |
| Disease | 377,250 | 275,044 | 27.1% | 11091886 |
| Anatomy | 39,362 | 33,877 | 13.9% | 11091803 |
| Protein Complex | 17,932 | 9,011 | 49.7% | 11091422 |
| Gene | 58,382,593 | 57,660,624 | 1.2% | 11092013 |
| Cell and Cell Line | 218,557 | 172,299 | 21.2% | 11091581 |

We found that semantic mapping assembly resulted in a meaningful consolidation of redundant terms within each domain (Reduction column in **Table 1**). Specifically, the high redundancy of resources in the domains of diseases and protein complexes lead to the largest reductions, especially given their relatively small size. Notably, the reduction for the gene domain is small because the number of model organism databases and the number of genes in each is only a small fraction of the NCBI Entrez Gene Database, which contains genes for several orders of magnitude more organisms. However, these estimations come with several caveats. First, these estimations can be artificially high because of missing mappings or due to the removal of many-to-many mappings during processing. Second, these estimations could be artificially low due to incorrect mappings or due to the missing terms from vocabularies that could not be accessed in full (e.g., SNOMED-CT for diseases).

Finally, we note that this analysis can be extended to other domains by writing new configurations for SeMRA. These could include other common entity types in biomedical research such as chemicals, protein families, pathways, molecular functions, assays, and taxa.

## 4 Discussion

We presented SeMRA, a configurable workflow for automatically assembling mappings at scale and an associated mapping database of 43.3 million mappings. SeMRA can be used to assemble context- and task-specific processed mappings using a combination of graph-based inference to infer indirect mappings, *a priori* domain knowledge, and custom prioritization that enables merging equivalent concepts.

The SeMRA mapping database is broader in scope compared to existing work on mapping integration. In comparison with SeMRA, [34] extracted 1 million database cross-references from a subset of the OBO Foundry comprising only 30 ontologies. The Ontology Mapping Service (OXO) imports most of the OBO Foundry and UMLS for supplementary mappings and contains 830 thousand mappings (as of 2024). TogoID and BridgeDB are relatively broader in coverage with 103 and 148 resources included, respectively. However, both BridgeDB and TogoID are limited in that they do not employ inference or make their results readily available in reusable formats.

### 4.1 Limitations

SeMRA’s reasoning approaches are inherently limited by the availability of mappings from the set of sources we integrated and the quality of such mappings. To mitigate this, the modular nature of SeMRA allows additional sources to be readily added by curating new prefixes and their associated ontology artifacts in the Bioregistry [27], by implementing additional database plugins in PyOBO [29], or by directly implementing custom processors in SeMRA. However, more generally, the limitations of existing mapping sources highlight the need to incorporate semi-automated mapping curation workflows like Biomappings [28] as well as automated approaches like Logmap [32], LOOM [15], or K-Boom [40].

The integration of mappings across fragmented sources is also challenging due to the fact that primary sources use inconsistent semantics when distributing mappings (for instance, as we observed in the case study on cell lines, there are database cross-references that result in many-to-many mappings). SeMRA attempts to mitigate this issue by (i) making available a combination of mutation features, (ii) configurability for the custom prioritization of resources across which mappings are traversed, and (iii) automated checking for inconsistencies induced by integrated mappings. Still, correctly resolving such inconsistencies is highly challenging and in some use cases, requires further custom logic beyond the starting point that SeMRA provides.

### 4.2 Future work on SeMRA and future vision

There are several open questions for developing methods for the inference and assembly of mappings. SeMRA does not yet explicitly handle alternative identifiers and replaced-by annotations in ontologies that describe when multiple terms cover the same concept. Further, splitting and merging terms creates additional challenges in reconciliation needed before making consistent mappings.

It would also be productive to explore how the construction of existing aggregate resources such as BridgeDB could be reproduced with a declarative configuration and automated pipeline using SeMRA. We expect this could significantly reduce the engineering effort needed to maintain such resources and increase the domains over which they are applicable.

We will also work toward improving the user experience for mapping curation in the SeMRA front-end in order to support and enable additional community curation. For example, this could be done by using SeMRA as a backend for feature-rich mapping curation interfaces like Cocoda [44]. However, irrespective of the interface, custom curation will require architectural support in SeMRA for integrating these during the generation of its mapping resource output which is done statically due to inference typically being a slow process necessitating.

SeMRA has the potential to be an important component of a broader ecosystem to actively monitor and support the maintenance of ontologies and databases. In this context, SeMRA could propose inferred mappings for curation by primary resource curators and alert them to inconsistencies (such as those caused by many-to-many mappings) on a timely basis. Such a service could be easily incorporated into resources that adhere to the open code, open data, open infrastructure (O3) guidelines [26], such as OBO Foundry ontologies. Ultimately, we envision SeMRA will act as an intermediate semantic interoperability layer allowing for the consistent usage of identifier resources in biomedicine and other fields.

## Acknowledgments

The authors thank Nicolas Matentzoglu for helpful discussions.

## Appendix

### Declarative Configuration

**Supplementary Figure S1:**
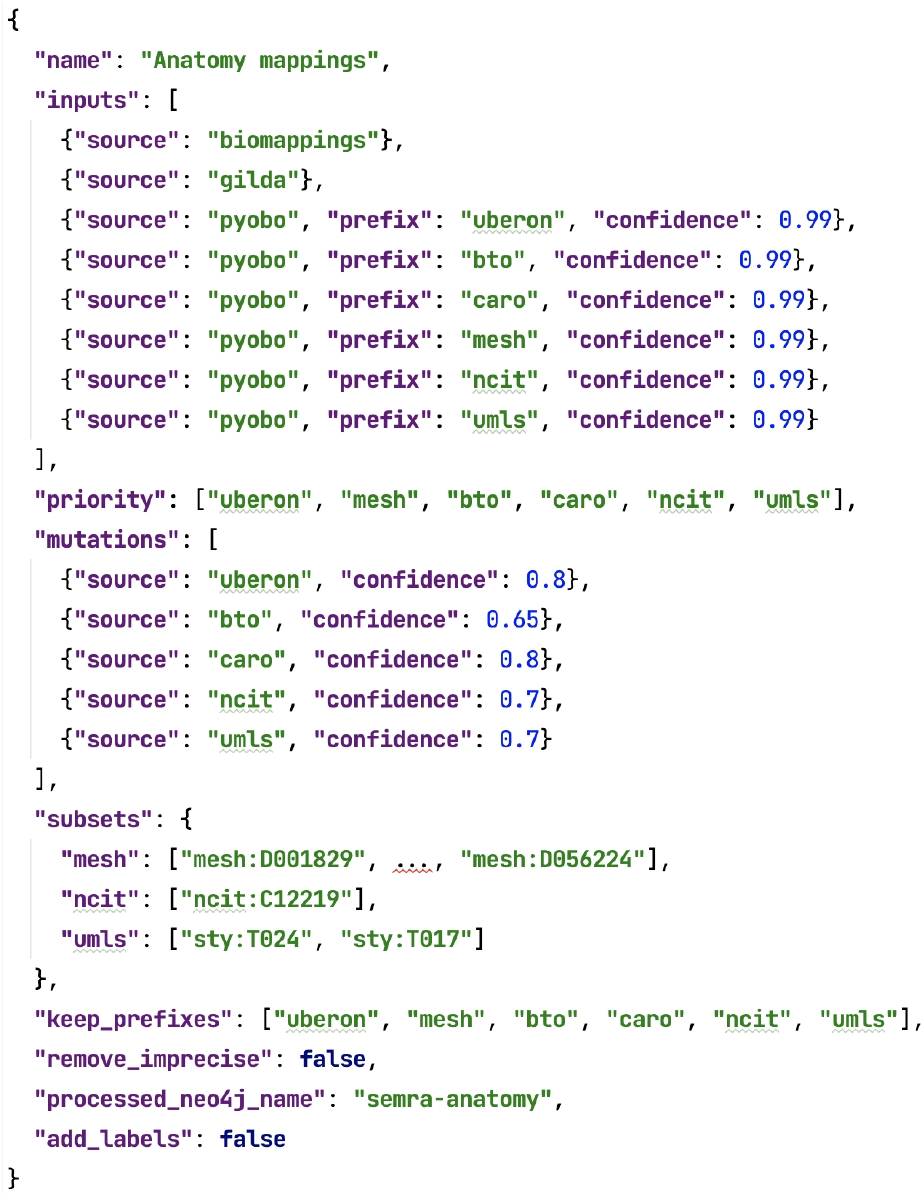
The declarative configuration for the Anatomy domain. MeSH subsets truncated for clarity.

The declarative configuration (**Supplementary Figure S1**) incorporates several fields:

1. *Name* a title for the configuration
2. *Inputs*, each containing a source API (e.g., PyOBO, Wikidata, Bioontologies, Custom), a prefix, and a consumerprovided confidence
3. *Subsets*, which allow for specifying when only an explicit list of sub-hierarchies in the resource should be used. This is only applicable for ontology-like resources
4. *Mutations*, a list of mutations objects. Commonly, this has a combination of a source and confidence, meaning that all database cross-references from the source can be implied to mean exact matches with the given confidence (besides many-to-many mappings, which are automatically removed)
5. *Keep Prefixes*, a list of prefixes to subset all mappings to before processing. *Post Keep Prefixes* does the same filter, but after processing. This is useful for incorporating resources like UMLS in the mapping process, but when it’s not desirable to retain mappings with UMLS entities due to licensing questions
6. *Priority*, the priority order for selecting a canonical entity to represent a clique of equivalent entities
7. *Remove Imprecise*, a flag for removing database cross-references that couldn’t be inferred as exact matches with adequate confidence
8. *Add Labels*, and other flags not shown, affect output by adding names to the SSSOM file. This can make processing take much longer as it requires caching all relevant external resources.
9. Not Shown: configuration for making outputs

### Web application

SeMRA provides a locally deployable web application to browse mappings, a snippet of which is shown in Supplementary Figure S2.

**Supplementary Figure S2:**
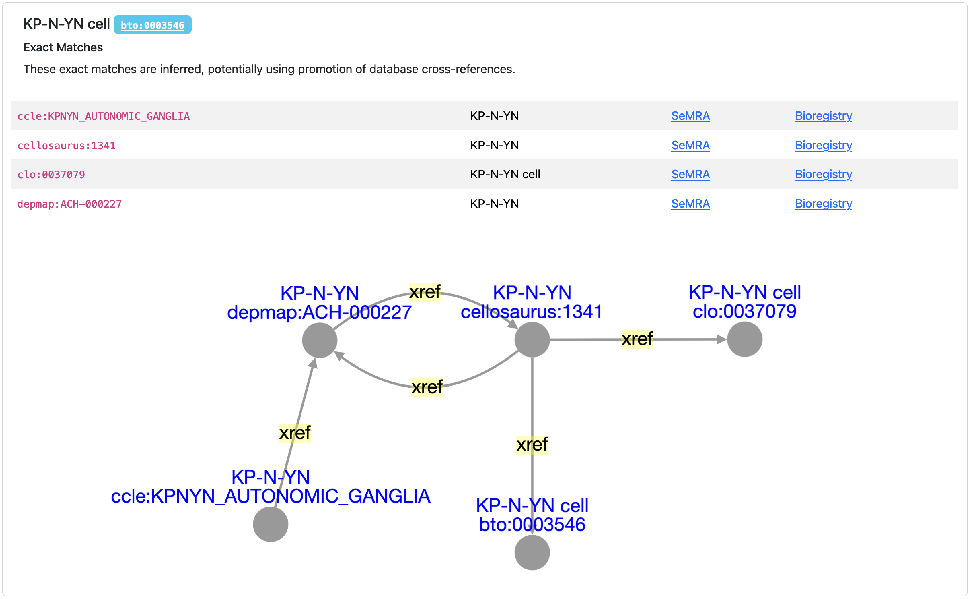
A view over the SeMRA web application.

### Mapping landscapes for specific entity types

**Supplementary Table S1:** Cell type and cell line landscape statistics showing the license for each resource, the version used to generate the results, the number of terms appearing in the landscape, and the scope for the inclusion of the resource: F is the full resource was used and S if a subset was used specific to cell types and cell lines.

| Resource | License | Version | Terms | Scope |
| --- | --- | --- | --- | --- |
| BTO [17] | CC-BY-4.0 | 2021-10-26 | 6,570 | F |
| CCLE [14] | ODbL-1.0 | 2019 | 1,061 | F |
| Cellosaurus [4] | CC-BY-4.0 | 51.0 | 159,463 | F |
| CL [10] | CC-BY-4.0 | 2025-02-13 | 3,046 | F |
| CLO [47] | CC-BY-3.0 | 2.1.188 | 39,125 | F |
| DepMap [5] | CC-BY-4.0 | 24Q4 | 1,814 | F |
| EFO [37] | Apache-2.0 | 3.76.0 | 27 | S |
| MeSH [45] | CC0-1.0 | 2025 | 636 | S |
| NCIT [16] | CC-BY-4.0 | 25.03c | 503 | S |
| UMLS [7] | Custom | 2024AB | 6,312 | S |

**Supplementary Table S2:** Disease landscape statistics, showing the license for each resource, the version used to generate the results, the number of terms appearing in the landscape, and the scope for the inclusion of the resource: F if the full resource was used, S if a subset specific to diseases, or O if only terms were included that were observed as mappings from other resources.

| Resource | License | Version | Terms | Scope |
| --- | --- | --- | --- | --- |
| DO [48] | CC0-1.0 | 2025-03-03 | 14,220 | F |
| EFO [37] | Apache-2.0 | 3.76.0 | 2,121 | S |
| GARD [24] | Unknown |  | 6,132 | F |
| ICD10 | Custom | 2019 | 2,345 | F |
| ICD10-CM | Custom |  | 2,830 | O |
| ICD11 | CC-BY-ND | 2025-01 | 71,175 | F |
| ICD9 | CC-BY-ND |  | 3,955 | O |
| ICD9-CM | CC-BY-ND |  | 2,226 | O |
| ICD-O | CC-BY-ND |  | 797 | O |
| MeSH [45] | CC0-1.0 | 2025 | 3,178 | S |
| MONDO [51] | CC-BY-4.0 | 2025-03-04 | 29,826 | F |
| NCIT [16] | CC-BY-4.0 | 25.03c | 20,522 | S |
| OMIM [1] | Custom | 2025-03-24 | 14,452 | O |
| OMIMPS [1] | Custom | 2025-03-24 | 588 | F |
| Orphanet [54] | CC-BY-4.0 | 4.6 | 15,507 | F |
| UMLS [7] | Custom | 2024AB | 187,376 | S |

## Footnotes

1 https://mapping-commons.github.io/sssom/chaining-rules

## References

[1] Joanna S. Amberger, Carol A. Bocchini, François Schiettecatte, Alan F. Scott, and Ada Hamosh. Omim.org: Online mendelian inheritance in man (omim®), an online catalog of human genes and genetic disorders. Nucleic Acids Research, 43:D789–D798, 1 2015. ISSN 1362-4962. doi: 10.1093/nar/gku1205.

[2] Alberto Anguita, Ana Escrich, and Victor Maojo. Fostering ontology alignment sharing: A general-purpose rdf mapping format. volume 192, page 970. IOS Press, 2013. ISBN 9781614992882. doi: 10.3233/978-1-61499-289-9-970.

[3] John A Bachman, Benjamin M Gyori, and Peter K Sorger. Automated assembly of molecular mechanisms at scale from text mining and curated databases. Molecular Systems Biology, 19, 5 2023. ISSN 1744-4292. doi: 10.15252/msb.202211325.

[4] Amos Bairoch. The Cellosaurus, a Cell-Line Knowledge Resource. J. Biomol. Tech., 29(2):25–38, jul 2018. ISSN 1943-4731 (Electronic). doi: 10.7171/jbt.18-2902-002.

[5] Jordi Barretina, Giordano Caponigro, Nicolas Stransky, Kavitha Venkatesan, Adam A Margolin, Sungjoon Kim, Christopher J Wilson, Joseph Lehàr, Gregory V Kryukov, Dmitriy Sonkin, Anupama Reddy, Manway Liu, Lauren Murray, Michael F Berger, John E Monahan, Paula Morais, Jodi Meltzer, Adam Korejwa, Judit Janè-Valbuena, Felipa A Mapa, Joseph Thibault, Eva Bric-Furlong, Pichai Raman, Aaron Shipway, Ingo H Engels, Jill Cheng, Guoying K Yu, Jianjun Yu, Peter Aspesi, Melanie de Silva, Kalpana Jagtap, Michael D Jones, Li Wang, Charles Hatton, Emanuele Palescandolo, Supriya Gupta, Scott Mahan, Carrie Sougnez, Robert C Onofrio, Ted Liefeld, Laura MacConaill, Wendy Winckler, Michael Reich, Nanxin Li, Jill P Mesirov, Stacey B Gabriel, Gad Getz, Kristin Ardlie, Vivien Chan, Vic E Myer, Barbara L Weber, Jeff Porter, Markus Warmuth, Peter Finan, Jennifer L Harris, Matthew Meyerson, Todd R Golub, Michael P Morrissey, William R Sellers, Robert Schlegel, and Levi A Garraway. The Cancer Cell Line Encyclopedia enables predictive modelling of anticancer drug sensitivity. Nature, 483(7391):603–607, 2012. ISSN 1476-4687. doi: 10.1038/nature11003. URL https://doi.org/10.1038/nature11003.

[6] Mark Birbeck and Shane McCarron. Curie syntax 1.0—a syntax for expressing compact uris. Working group note, W3C, 2010.

[7] O. Bodenreider. The unified medical language system (umls): integrating biomedical terminology. Nucleic Acids Research, 32:267D–270, 1 2004. ISSN 1362-4962. doi: 10.1093/nar/gkh061.

[8] Mapping Commons. Sssom-py: A python library for working with simple standard for sharing ontological mappings (sssom). https://github.com/mapping-commons/sssom-py, 2024.

[9] Jèròme David, Jèròme Euzenat, François Scharffe, and Càssia Trojahn dos Santos. The alignment api 4.0. Semant. Web, 2(1):3–10, jan 2011. ISSN 1570-0844.

[10] Alexander D Diehl, Terrence F Meehan, Yvonne M Bradford, Matthew H Brush, Wasila M Dahdul, David S Dougall, Yongqun He, David Osumi-Sutherland, Alan Ruttenberg, Sirarat Sarntivijai, Ceri E Van Slyke, Nicole A Vasilevsky, Melissa A Haendel, Judith A Blake, and Christopher J Mungall. The Cell Ontology 2016: enhanced content, modularization, and ontology interoperability. J. Biomed. Semantics, 7(1):44, 2016. ISSN 2041-1480. doi: 10.1186/s13326-016-0088-7. URL https://doi.org/10.1186/s13326-016-0088-7.

[11] Marcel Friedrichs. Biodwh2: an automated graph-based data warehouse and mapping tool. Journal of Integrative Bioinformatics, 18(2):167–176, 2021. doi: 10.1515/jib-2020-0033. URL https://doi.org/10.1515/jib-2020-0033.

[12] Anna Gaulton, Anne Hersey, Micha L. Nowotka, A. Patricia Bento, Jon Chambers, David Mendez, Prudence Mutowo, Francis Atkinson, Louisa J. Bellis, Elena Cibrian-Uhalte, Mark Davies, Nathan Dedman, Anneli Karlsson, Mària Paula Magarinos, John P. Overington, George Papadatos, Ines Smit, and Andrew R. Leach. The ChEMBL database in 2017. Nucleic Acids Res., 45(D1):D945–D954, 2017. ISSN 13624962. doi: 10.1093/nar/gkw1074.

[13] David Geleta, Andriy Nikolov, Mark ODonoghue, Benedek Rozemberczki, Anna Gogleva, Valentina Tamma, and Terry R. Payne. Ontomerger: An ontology integration library for deduplicating and connecting knowledge graph nodes. arXiv, 2022. doi: 10.48550/ARXIV.2206.02238. URL https://arxiv.org/abs/2206.02238.

[14] Mahmoud Ghandi, Franklin W Huang, Judit Janè-Valbuena, Gregory V Kryukov, Christopher C Lo, E Robert McDonald, Jordi Barretina, Ellen T Gelfand, Craig M Bielski, Haoxin Li, Kevin Hu, Alexander Y Andreev-Drakhlin, Jaegil Kim, Julian M Hess, Brian J Haas, François Aguet, Barbara A Weir, Michael V Rothberg, Brenton R Paolella, Michael S Lawrence, Rehan Akbani, Yiling Lu, Hong L Tiv, Prafulla C Gokhale, Antoine de Weck, Ali Amin Mansour, Coyin Oh, Juliann Shih, Kevin Hadi, Yanay Rosen, Jonathan Bistline, Kavitha Venkatesan, Anupama Reddy, Dmitriy Sonkin, Manway Liu, Joseph Lehar, Joshua M Korn, Dale A Porter, Michael D Jones, Javad Golji, Giordano Caponigro, Jordan E Taylor, Caitlin M Dunning, Amanda L Creech, Allison C Warren, James M McFarland, Mahdi Zamanighomi, Audrey Kauffmann, Nicolas Stransky, Marcin Imielinski, Yosef E Maruvka, Andrew D Cherniack, Aviad Tsherniak, Francisca Vazquez, Jacob D Jaffe, Andrew A Lane, David M Weinstock, Cory M Johannessen, Michael P Morrissey, Frank Stegmeier, Robert Schlegel, William C Hahn, Gad Getz, Gordon B Mills, Jesse S Boehm, Todd R Golub, Levi A Garraway, and William R Sellers. Next-generation characterization of the Cancer Cell Line Encyclopedia. Nature, 569(7757):503–508, 2019. ISSN 1476-4687. doi: 10.1038/s41586-019-1186-3. URL https://doi.org/10.1038/s41586-019-1186-3.

[15] Amir Ghazvinian, Natalya F Noy, and Mark A Musen. Creating mappings for ontologies in biomedicine: simple methods work. AMIA … Annu. Symp. proceedings. AMIA Symp., 2009:198–202, nov 2009. ISSN 1942-597X (Electronic).

[16] Jennifer Golbeck, Gilberto Fragoso, Frank Hartel, Jim Hendler, Jim Oberthaler, and Bijan Parsia. The national cancer institute’s thèsaurus and ontology. Journal of Web Semantics, 1:75–80, 12 2003. ISSN 15708268. doi: 10.1016/j.websem.2003.07.007.

[17] Marion Gremse, Antje Chang, Ida Schomburg, Andreas Grote, Maurice Scheer, Christian Ebeling, and Dietmar Schomburg. The BRENDA Tissue Ontology (BTO): the first all-integrating ontology of all organisms for enzyme sources. Nucleic Acids Res., 39(Suppl 1):D507–D513, jan 2011. ISSN 0305-1048. doi: 10.1093/nar/gkq968. URL https://doi.org/10.1093/nar/gkq968.

[18] Xiuzhan Guo, Arthur Berrill, Ajinkya Kulkarni, Kostya Belezko, and Min Luo. Merging ontologies algebraically. arXiv, 2022. doi: 10.48550/ARXIV.2208.08715. URL https://arxiv.org/abs/2208.08715.

[19] Benjamin M Gyori, John A Bachman, Kartik Subramanian, Jeremy L Muhlich, Lucian Galescu, and Peter K Sorger. From word models to executable models of signaling networks using automated assembly. Mol. Syst. Biol., 13(11):954, 2017. ISSN 1744-4292. doi: 10.15252/msb.20177651. URL http://msb.embopress.org/lookup/doi/10.15252/msb.20177651.

[20] Benjamin M Gyori, Charles Tapley Hoyt, and Albert Steppi. Gilda: biomedical entity text normalization with machine-learned disambiguation as a service. Bioinformatics Advances, 2(1), 05 2022. ISSN 2635-0041. doi: 10.1093/bioadv/vbac034. URL https://doi.org/10.1093/bioadv/vbac034.vbac034.

[21] Melissa Haendel, Nicole Vasilevsky, Deepak Unni, Cristian Bologa, Nomi Harris, Heidi Rehm, Ada Hamosh, Gareth Baynam, Tudor Groza, Julie McMurry, Hugh Dawkins, Ana Rath, Courtney Thaxton, Giovanni Bocci, Marcin P. Joachimiak, Sebastian Köhler, Peter N. Robinson, Chris Mungall, and Tudor I. Oprea. How many rare diseases are there? Nature Reviews Drug Discovery, 19:77–78, 2 2020. ISSN 1474-1776. doi: 10.1038/d41573-019-00180-y.

[22] Ian Harrow, Rama Balakrishnan, Ernesto Jimenez-Ruiz, Simon Jupp, Jane Lomax, Jane Reed, Martin Romacker, Christian Senger, Andrea Splendiani, Jabe Wilson, and Peter Woollard. Ontology mapping for semantically enabled applications. Drug Discovery Today, 24:2068–2075, 10 2019. ISSN 13596446. doi: 10.1016/j.drudis.2019.05.020.

[23] Daniel Scott Himmelstein, Antoine Lizee, Christine Hessler, Leo Brueggeman, Sabrina L Chen, Dexter Hadley, Ari Green, Pouya Khankhanian, and Sergio E Baranzini. Systematic integration of biomedical knowledge prioritizes drugs for repurposing. Elife, 6, sep 2017. ISSN 2050-084X. doi: 10.7554/eLife.26726. URL http://www.ncbi.nlm.nih.gov/pubmed/28936969 http://www.pubmedcentral.nih.gov/articlerender.fcgi?artid=PMC5640425.

[24] Amber Hoskins. Genetic and rare diseases information center (gard). Medical reference services quarterly, 41:389–394, 2022. ISSN 1540-9597. doi: 10.1080/02763869.2022.2131143.

[25] Charles Tapley Hoyt. biopragmatics/bioontologies: v0.7.0, March 2025. URL https://doi.org/10.5281/zenodo.15075084.

[26] Charles Tapley Hoyt and Benjamin M Gyori. The O3 guidelines: open data, open code, and open infrastructure for sustainable curated scientific resources. Scientific Data, 11(1):547, 2024.

[27] Charles Tapley Hoyt, Meghan Balk, Tiffany J Callahan, Daniel Domingo-Fernàndez, Melissa A Haendel, Harshad B Hegde, Daniel S Himmelstein, Klas Karis, John Kunze, Tiago Lubiana, Nicolas Matentzoglu, Julie McMurry, Sierra Moxon, Christopher J Mungall, Adriano Rutz, Deepak R Unni, Egon Willighagen, Donald Winston, and Benjamin M Gyori. Unifying the identification of biomedical entities with the Bioregistry. Sci. Data, 9(1):714, 2022. ISSN 2052-4463. doi: 10.1038/s41597-022-01807-3. URL https://doi.org/10.1038/s41597-022-01807-3.

[28] Charles Tapley Hoyt, Amelia L Hoyt, and Benjamin M Gyori. Prediction and curation of missing biomedical identifier mappings with biomappings. Bioinformatics, 39, 4 2023. ISSN 1367-4803. doi: 10.1093/bioinformatics/btad130.

[29] Charles Tapley Hoyt, Benjamin M. Gyori, Daniel Domingo-Fernàndez, Tenzin Nanglo, Harshad, Sam, Jose P. Faria, and Klas Karis. biopragmatics/pyobo: v0.11.2, December 2024. URL 10.5281/zenodo.14501873.

[30] Shuya Ikeda, Hiromasa Ono, Tazro Ohta, Hirokazu Chiba, Yuki Naito, Yuki Moriya, Shuichi Kawashima, Yasunori Yamamoto, Shinobu Okamoto, Susumu Goto, and Toshiaki Katayama. TogoID: an exploratory ID converter to bridge biological datasets. Bioinformatics, 07 2022. ISSN 1367-4803. doi: 10.1093/bioinformatics/btac491. URL https://doi.org/10.1093/bioinformatics/btac491.btac491.

[31] Rebecca C. Jackson, Nicolas Matentzoglu, James A. Overton, Randi Vita, James P. Balhoff, Pier Luigi Buttigieg, Seth Carbon, Melanie Courtot, Alexander D. Diehl, Damion Dooley, William Duncan, Nomi L. Harris, Melissa A. Haendel, Suzanna E. Lewis, Darren A. Natale, David Osumi-Sutherland, Alan Ruttenberg, Lynn M. Schriml, Barry Smith, Christian J. Stoeckert, Nicole A. Vasilevsky, Ramona L. Walls, Jie Zheng, Christopher J. Mungall, and Bjoern Peters. OBO Foundry in 2021: Operationalizing Open Data Principles to Evaluate Ontologies. Database (Oxford)., 2021(October):1–9, 2021. ISSN 1758-0463. doi: 10.1093/database/baab069.

[32] Ernesto Jimènez-Ruiz and Bernardo Cuenca Grau. LogMap: Logic-Based and Scalable Ontology Matching. In Lora Aroyo, Chris Welty, Harith Alani, Jamie Taylor, Abraham Bernstein, Lalana Kagal, Natasha Noy, and Eva Blomqvist, editors, The Semantic Web - ISWC 2011, pages 273–288, Berlin, Heidelberg, 2011. Springer Berlin Heidelberg. ISBN 978-3-642-25073-6.

[33] Max Karlsson, Cheng Zhang, Loren Meár, Wen Zhong, Andreas Digre, Borbala Katona, Evelina Sjöstedt, Lynn Butler, Jacob Odeberg, Philip Dusart, Fredrik Edfors, Per Oksvold, Kalle von Feilitzen, Martin Zwahlen, Muhammad Arif, Ozlem Altay, Xiangyu Li, Mehmet Ozcan, Adil Mardinoglu, Linn Fagerberg, Jan Mulder, Yonglun Luo, Fredrik Ponten, Mathias Uhlèn, and Cecilia Lindskog. A single–cell type transcriptomics map of human tissues. Science Advances, 7, 7 2021. ISSN 2375-2548. doi: 10.1126/sciadv.abh2169.

[34] Amir Laadhar, Élcio Abrahão, and Clèment Jonquet. Investigating one million xrefs in thirthy ontologies from the OBO world. In Janna Hastings and Frank Loebe, editors, ICBO 2020, volume 2807 of CEUR Workshop Proceedings, pages 1–12. CEUR-WS.org, 2020. URL http://ceur-ws.org/Vol-2807/paperG.pdf.

[35] Patrick Lambrix and He Tan. Ontology Alignment and Merging, pages 133–149. Springer London, 2008. ISBN 978-1-84628-885-2. doi: 10.1007/978-1-84628-885-26. URL https://doi.org/10.1007/978-1-84628-885-2_6.

[36] Donna Maglott, Jim Ostell, Kim D. Pruitt, and Tatiana Tatusova. Entrez gene: Gene-centered information at NCBI. Nucleic Acids Res., 39(SUPPL. 1):52–57, 2011. ISSN 03051048. doi: 10.1093/nar/gkq1237.

[37] James Malone, Ele Holloway, Tomasz Adamusiak, Misha Kapushesky, Jie Zheng, Nikolay Kolesnikov, Anna Zhukova, Alvis Brazma, and Helen Parkinson. Modeling sample variables with an Experimental Factor Ontology. Bioinformatics, 26(8):1112–1118, 03 2010. ISSN 1367-4803. doi: 10.1093/bioinformatics/btq099. URL https://doi.org/10.1093/bioinformatics/btq099.

[38] Nicolas Matentzoglu, James P Balhoff, Susan M Bello, Chris Bizon, Matthew Brush, Tiffany J Callahan, Christopher G Chute, William D Duncan, Chris T Evelo, Davera Gabriel, John Graybeal, Alasdair Gray, Benjamin M Gyori, Melissa Haendel, Henriette Harmse, Nomi L Harris, Ian Harrow, Harshad B Hegde, Amelia L Hoyt, Charles T Hoyt, Dazhi Jiao, Ernesto Jimènez-Ruiz, Simon Jupp, Hyeongsik Kim, Sebastian Koehler, Thomas Liener, Qinqin Long, James Malone, James A McLaughlin, Julie A McMurry, Sierra Moxon, Monica C Munoz-Torres, David Osumi-Sutherland, James A Overton, Bjoern Peters, Tim Putman, Núria Queralt-Rosinach, Kent Shefchek, Harold Solbrig, Anne Thessen, Tania Tudorache, Nicole Vasilevsky, Alex H Wagner, and Christopher J Mungall. A Simple Standard for Sharing Ontological Mappings (SSSOM). Database, 2022:baac035, jan 2022. ISSN 1758-0463. doi: 10.1093/database/baac035. URL https://doi.org/10.1093/database/baac035.

[39] Nicolas Matentzoglu, Joe Flack, John Graybeal, Nomi L. Harris, Harshad B. Hegde, Charles T. Hoyt, Hyeongsik Kim, Sabrina Toro, Nicole Vasilevsky, and Christopher J. Mungall. A simple standard for ontological mappings 2022: Updates of data model and outlook. CEUR Workshop Proceedings, 3324:61–66, 2022. ISSN 1613-0073.

[40] Christopher J. Mungall, Sebastian Koehler, Peter Robinson, Ian Holmes, and Melissa Haendel. k-boom: A bayesian approach to ontology structure inference, with applications in disease ontology construction. bioRxiv, 2019. doi: 10.1101/048843. URL https://www.biorxiv.org/content/early/2019/01/29/048843.

[41] David N. Nicholson and Casey S. Greene. Constructing knowledge graphs and their biomedical applications. Comput. Struct. Biotechnol. J., 18:1414–1428, 2020. ISSN 20010370. doi: 10.1016/j.csbj.2020.05.017. URL https://doi.org/10.1016/j.csbj.2020.05.017.

[42] Natalya F Noy, Sherri de Coronado, Harold Solbrig, Gilberto Fragoso, Frank W Hartel, and Mark A Musen. Representing the nci thesaurus in owl dl: Modeling tools help modeling languages. Applied ontology, 3:173– 190, 1 2008. ISSN 1570-5838. doi: 10.3233/AO-2008-0051.

[43] Dexter Pratt, Jing Chen, David Welker, Ricardo Rivas, Rudolf Pillich, Vladimir Rynkov, Keiichiro Ono, Carol Miello, Lyndon Hicks, Sandor Szalma, Aleksandar Stojmirovic, Radu Dobrin, Michael Braxenthaler, Jan Kuentzer, Barry Demchak, and Trey Ideker. NDEx, the Network Data Exchange. Cell Syst., 1(4):302– 305, 2015. ISSN 24054712. doi: 10.1016/j.cels.2015.10.001. URL http://dx.doi.org/10.1016/j.cels.2015.10.001.

[44] Fernanda Ribeiro and Maria Elisa Cerveira, editors. Challenges and Opportunities for Knowledge Organization in the Digital Age, volume 16 of Advances in Knowledge Organization. Ergon, Baden-Baden, 1 edition, 2018.

[45] FB Rogers. Medical subject headings. Bull. Med. Libr. Assoc., 51:114–6, jan 1963. ISSN 0025-7338. URL http://www.ncbi.nlm.nih.gov/pubmed/13982385http://www.pubmedcentral.nih.gov/articlerender.fcgi?artid=PMC197951.

[46] P. Romano, A. Manniello, O. Aresu, M. Armento, M. Cesaro, and B. Parodi. Cell line data base: structure and recent improvements towards molecular authentication of human cell lines. Nucleic Acids Research, 37:D925–D932, 1 2009. ISSN 0305-1048. doi: 10.1093/nar/gkn730.

[47] Sirarat Sarntivijai, Yu Lin, Zuoshuang Xiang, Terrence F Meehan, Alexander D Diehl, Uma D Vempati, Stephan C Schürer, Chao Pang, James Malone, Helen Parkinson, Yue Liu, Terue Takatsuki, Kaoru Saijo, Hiroshi Masuya, Yukio Nakamura, Matthew H Brush, Melissa A Haendel, Jie Zheng, Christian J Stoeckert, Bjoern Peters, Christopher J Mungall, Thomas E Carey, David J States, Brian D Athey, and Yongqun He. CLO: The cell line ontology. J. Biomed. Semantics, 5(1):37, 2014. ISSN 2041-1480. doi: 10.1186/2041-1480-5-37. URL https://doi.org/10.1186/2041-1480-5-37.

[48] Lynn M Schriml, James B Munro, Mike Schor, Dustin Olley, Carrie McCracken, Victor Felix, J Allen Baron, Rebecca Jackson, Susan M Bello, Cynthia Bearer, Richard Lichenstein, Katharine Bisordi, Nicole Campion Dialo, Michelle Giglio, and Carol Greene. The human disease ontology 2022 update. Nucleic Acids Research, 50(D1):D1255–D1261, 11 2021. ISSN 0305-1048. doi: 10.1093/nar/gkab1063. URL https://doi.org/10.1093/nar/gkab1063.

[49] Aviad Tsherniak, Francisca Vazquez, Phil G Montgomery, Barbara A Weir, Gregory Kryukov, Glenn S Cowley, Stanley Gill, William F Harrington, Sasha Pantel, John M Krill-Burger, et al. Defining a cancer dependency map. Cell, 170(3):564–576, 2017.

[50] Martijn P. van Iersel, Alexander R. Pico, Thomas Kelder, Jianjiong Gao, Isaac Ho, Kristina Hanspers, Bruce R. Conklin, and Chris T. Evelo. The BridgeDb framework: Standardized access to gene, protein and metabolite identifier mapping services. BMC Bioinformatics, 11, 2010. ISSN 14712105. doi: 10.1186/1471-2105-11-5.

[51] Nicole A Vasilevsky, Nicolas A Matentzoglu, Sabrina Toro, Joseph E Flack, Harshad Hegde, Deepak R Unni, Gioconda F Alyea, Joanna S Amberger, Larry Babb, James P Balhoff, Taylor I Bingaman, Gully A Burns, Orion J Buske, Tiffany J Callahan, Leigh C Carmody, Paula Carrio Cordo, Lauren E Chan, George S Chang, Sean L Christiaens, Louise C Daugherty, Michel Dumontier, Laura E Failla, May J Flowers, H Alpha Garrett, Jennifer L Goldstein, Dylan Gration, Tudor Groza, Marc Hanauer, Nomi L Harris, Jason A Hilton, Daniel S Himmelstein, Charles Tapley Hoyt, Megan S Kane, Sebastian Köhler, David Lagorce, Abbe Lai, Martin Larralde, Antonia Lock, Irene López Santiago, Donna R Maglott, Adriana J Malheiro, Birgit H M Meldal, Monica C Munoz-Torres, Tristan H Nelson, Frank W Nicholas, David Ochoa, Daniel P Olson, Tudor I Oprea, David Osumi-Sutherland, Helen Parkinson, Zoë May Pendlington, Ana Rath, Heidi L Rehm, Lyubov Remennik, Erin R Riggs, Paola Roncaglia, Justyne E Ross, Marion F Shadbolt, Kent A Shefchek, Morgan N Similuk, Nicholas Sioutos, Damian Smedley, Rachel Sparks, Ray Stefancsik, Ralf Stephan, Andrea L Storm, Doron Stupp, Gregory S Stupp, Jagadish Chandrabose Sundaramurthi, Imke Tammen, Darin Tay, Courtney L Thaxton, Eloise Valasek, Jordi Valls-Margarit, Alex H Wagner, Danielle Welter, Patricia L Whetzel, Lori L Whiteman, Valerie Wood, Colleen H Xu, Andreas Zankl, Xingmin Aaron Zhang, Christopher G Chute, Peter N Robinson, Christopher J Mungall, Ada Hamosh, and Melissa A Haendel. Mondo: Unifying diseases for the world, by the world. medRxiv, 2022. doi: 10.1101/2022.04.13.22273750. URL https://www.medrxiv.org/content/early/2022/05/03/2022.04.13.22273750.

[52] Uma D. Vempati, Caty Chung, Chris Mader, Amar Koleti, Nakul Datar, Dušica Vidović, David Wrobel, Sean Erickson, Jeremy L. Muhlich, Gabriel Berriz, Cyril H. Benes, Aravind Subramanian, Ajay Pillai, Caroline E. Shamu, and Stephan C. Schürer. Metadata standard and data exchange specifications to describe, model, and integrate complex and diverse high-throughput screening data from the library of integrated network-based cellular signatures (lincs). SLAS Discovery, 19:803–816, 6 2014. ISSN 24725552. doi: 10.1177/1087057114522514.

[53] Denny Vrandečić and Markus Krötzsch. Wikidata: a free collaborative knowledgebase. Communications of the ACM, 57(10):78–85, 2014.

[54] SS Weinreich, R Mangon, JJ Sikkens, ME en Teeuw, and MC Cornel. [orphanet: a european database for rare diseases]. Nederlands tijdschrift voor geneeskunde, 152:518–9, 3 2008. ISSN 0028-2162.

[55] Patricia L Whetzel, Natalya F Noy, Nigam H Shah, Paul R Alexander, Csongor Nyulas, Tania Tudorache, and Mark A Musen. BioPortal: enhanced functionality via new Web services from the National Center for Biomedical Ontology to access and use ontologies in software applications. Nucleic Acids Res., 39(Web Server issue):W541–5, jul 2011. ISSN 1362-4962 (Electronic). doi: 10.1093/nar/gkr469.

[56] Mark D Wilkinson, Michel Dumontier, IJsbrand Jan Aalbersberg, Gabrielle Appleton, Myles Axton, Arie Baak, Niklas Blomberg, Jan-Willem Boiten, Luiz Bonino da Silva Santos, Philip E Bourne, Jildau Bouwman, Anthony J Brookes, Tim Clark, Mercé Crosas Ingrid Dillo, Olivier Dumon, Scott Edmunds, Chris T Evelo, Richard Finkers, Alejandra Gonzalez-Beltran, Alasdair J G Gray, Paul Groth, Carole Goble, Jeffrey S Grethe, Jaap Heringa, Peter AC ‘t Hoen, Rob Hooft, Tobias Kuhn, Ruben Kok, Joost Kok, Scott J Lusher, Maryann E Martone, Albert Mons, Abel L Packer, Bengt Persson, Philippe Rocca-Serra, Marco Roos, Rene van Schaik, Susanna-Assunta Sansone, Erik Schultes, Thierry Sengstag, Ted Slater, George Strawn, Morris A Swertz, Mark Thompson, Johan van der Lei, Erik van Mulligen, Jan Velterop, Andra Waagmeester, Peter Wittenburg, Katherine Wolstencroft, Jun Zhao, and Barend Mons. The FAIR Guiding Principles for scientific data management and stewardship. Sci. Data, 3(1):160018, 2016. ISSN 2052-4463. doi: 10.1038/sdata.2016.18. URL https://doi.org/10.1038/sdata.2016.18.

